# A chromosome-level genome assembly of the Eurasian great grey owl, *Strix nebulosa lapponica* (Thunberg 1798)

**DOI:** 10.64898/2026.06.29.735218

**Authors:** Marius A. Strand, Inga A. Frøland Steindal, Erlend Ragnhildstveit, Roar Solheim, Ole K. Tørresen, Morten Skage, Giada Ferrari, Ave Tooming-Klunderud, Kjetill S. Jakobsen

## Abstract

We present a chromosome-level genome assembly of a female great grey owl (*Strix nebulosa lapponica*). The assembly comprises two pseudo-haplotypes of 1554 Mb and 1242 Mb, with 83.2% and 91.4% scaffolded into 40 autosomal chromosomes, in addition to the W and Z sex chromosomes both placed in hap1. Assembly completeness is high (BUSCO 99.2% and 94.8%), with 18,493 and 17,279 annotated protein-coding genes for hap1 and hap2, respectively. This genome establishes a reference for investigating genetic variation and chromosome evolution in great grey owls. Compared with the previous *S. nebulosa* assembly, this assembly includes both sex chromosomes, separates regions that were previously collapsed, and resolves 82 chromosomes total. While larger chromosomes show broadly conserved synteny across owl assemblies, the recovery of additional conserved microchromosome-associated genes suggests that ONT reads improved resolution of the smallest chromosomes relative to HiFi-based assemblies.

## Introduction

The great grey owl (*Strix nebulosa lapponica*) is among the largest owl species in the Northern Hemisphere, with a Holarctic distribution spanning boreal and subarctic regions of both Eurasia and North America (Cramp, 1985; del Hoyo et al., 1999). Two subspecies are recognized: *Strix nebulosa lapponica* in Eurasia and S*trix nebulosa nebulosa* in North America (del Hoyo et al., 1999; Mikkola, 1983). These subspecies are broadly similar in size, morphology, and behaviour, differing mainly in plumage characteristics, with slight variation in coloration and patterning, while ecological traits remain largely conserved across continents (Cramp, 1985; del Hoyo et al., 1999; Solheim, 2020).

Great grey owls have a body length typically ranging from approximately 60 to 70 cm and a wingspan exceeding 140 cm (del Hoyo et al., 1999; Mikkola, 1983). However, they have relatively low body mass compared to other large raptors, generally weighing between 800 and 1500 g (Mikkola, 1983). The distinct wing bar patterns, along with moult characteristics, have been demonstrated to vary with age and can be used for individual identification, ageing and demographic studies (Solheim, 2016, 2020).

The species primarily inhabits boreal coniferous forests and forest edges, often associated with open areas such as clear-cuts, bogs, and meadows (Cramp, 1985; del Hoyo et al., 1999). Great grey owls are typically crepuscular hunters but may also be active during daylight (Mikkola, 1983, Stefansson, 1997). Their hunting behaviour is highly specialized, relying extensively on auditory cues to detect microtines beneath vegetation or snow. Individuals frequently hunt from low perches, such as tree stumps or fence posts, and may glide or hover before executing a characteristic plunge through snow or ground cover to capture prey (Clark et al., 2022).

Great grey owls are not nestbuilders, but use twignests built by raptors like goshawks (*Accipiter gentilis*) or buzzards (*Buteo* sp.), and occasionally ravens (*Corvus corax*), typically located in mature forest stands (Cramp, 1985; Stefansson, 1997). Clutch size varies from 2 to 5 eggs, although up to 7 eggs may be laid in years of high microtine abundance (Cramp, 1985; Mikkola, 1983). Breeding frequency and success is strongly influenced by temporal fluctuations in food resources, and the owls are particularly dependent on microtines under 50g. Voles and wood lemmings (*Myopus schisticolor*) have historically had cycling population numbers with peaks on average every 3-4 years (Angerbjörn et al., 2001; Wilson et al., 2017). From late 1980-ies to ca 2005 microtine population cycles appeared to level off in large parts of Fennoscandia, with low peak years and especially *Microtus* species largely absent. Changing climate was believed to be a main cause of these abbreviations, but cycles and population peaks regained their former strength after 2005 (Solheim, 2020).

The great grey owl exhibits flexible movement patterns. While often associated with relatively stable home ranges, individuals may undertake substantial dispersal or irruptive movements in response to changing environmental conditions (Newton, 2008; Solheim et al. in prep.). These movements may span over a thousand kilometres between breeding seasons (Solheim & Stefansson, 2016; Valkama et al., 2014; Solheim et al. in prep.) and contribute to shifts in local population structure and distribution. The great grey owl has expanded its breeding range in Scandinavia in a south-southwesterly direction over the last 25 years (Ottosson et al., 2025). Particularly interesting is the expansion into southeastern Norway of birds originating from the Swedish population: breeding pairs have rapidly increased in this region since the early 2000s, with 140 nests recorded in 2025 in Hedmark county (Solheim & Berg, 2025). During the same period, there have also been expansions of ssp. *lapponica* in a south-westerly direction from Russia into Poland, Belarus and the Baltics (Ławicki et al., 2013). In light of this ongoing range expansion, long-distance movements, and the strong influence of fluctuating prey resources on reproduction, a high-quality, chromosome-resolved genome of the ssp. *lapponica*, is crucial for future studies of population dynamics and demographic history across the species’ changing range.

Here, we present a high quality chromosome-resolved assembly of the Eurasian great grey owl (*S. nebulosa lapponica*), generated by long-read Oxford Nanopore and Hi-C sequencing. It will serve as a resource for comparative genomic research, evolution of owls and local adaptation in great grey owls. Furthermore, it will be a crucial resource for studying population genetic structure and dynamics, helping us understand and monitor the species amid rapid changes in its distribution and breeding grounds.

## Material and Methods

### Sample acquisition and DNA extraction

A blood sample from a female *Strix nebulosa lapponica* specimen (“Siv”) was collected from Nord-Odal, Innlandet county, Norway (60°28’15.1”N 11°26’37.9”E) on 5^th^ of June 2025. The area is not a protected nature reserve. The female was caught with a net next to the nesting platform (50×50 cm area with a 15 cm frame, filled with approximately 10 cm layer of wood chips from decaying trees, 5 metres above ground in a large pine), located in a biotope dominated by sparsely distributed Scots pine trees (*Pinus sylvestris*) and 50 metres from a bog (Figure 1C). There were originally 4 eggs in the nest, but 2 chicks, approximately four weeks of age, when the owl was sampled. She was observed visiting the platform again shortly after she was released. To reduce time of handling no measurements or weight data was collected from the female. The female was first ringed as an adult (4 years or older) breeding bird in 2021, nesting 16 km further south in Eidsvoll municipality, so at least 8 years or older when reencountered in 2025. A blood sample was taken from the basilic vein of the wing and stored in an insulated ice box for approximately 4 hours before freezing at −80 °C in the laboratory. Blood sample was taken under FOTS 31029 from the Norwegian animal ethics authority Mattilsynet.

**Figure 1.**
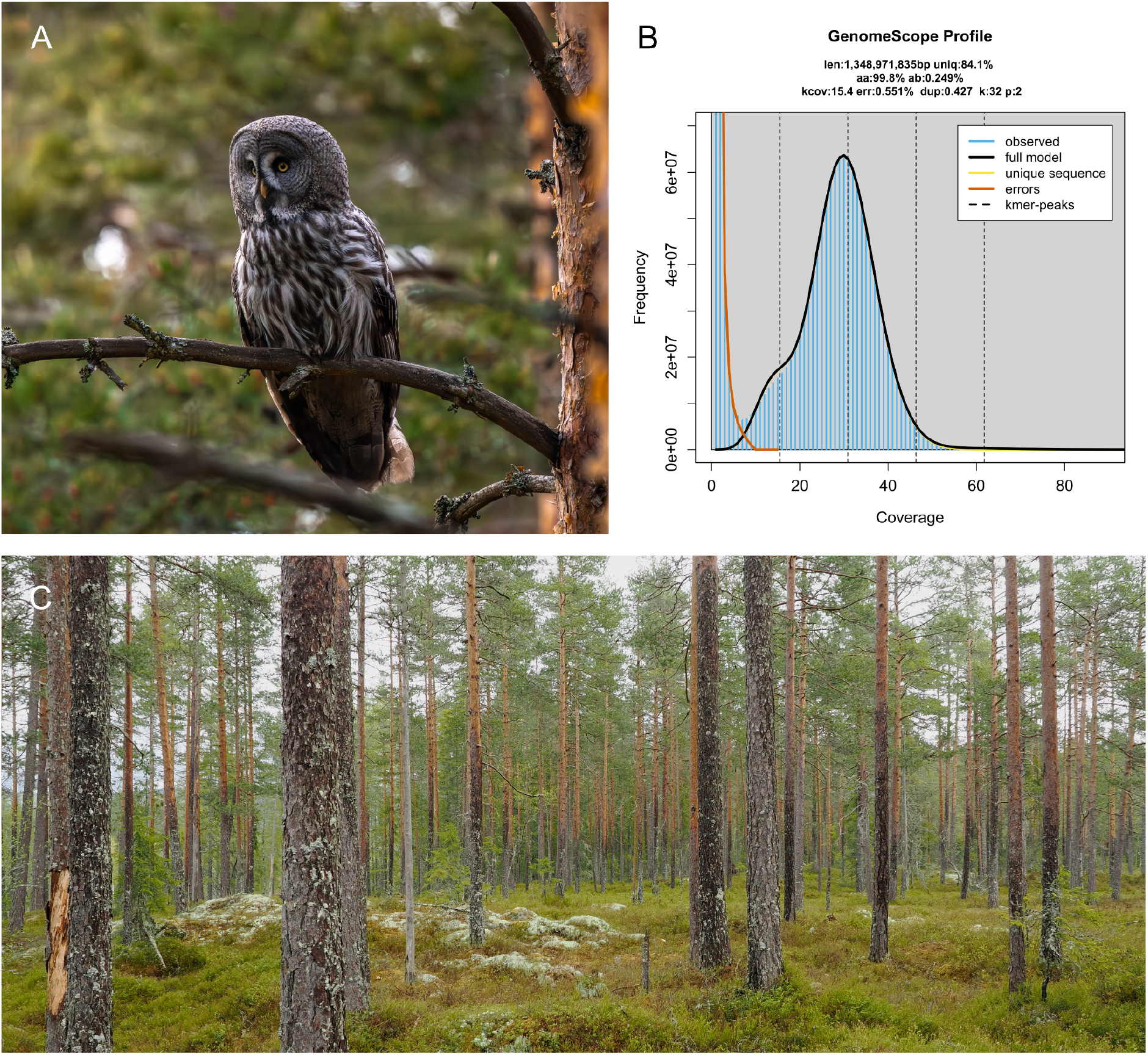
Representative specimen and genome profile. **A)** Photo of a representative female specimen from the sampling area; photo by Inga A. Frøland Steindal. **B)** GenomeScope profile of the ONT reads from the sequenced individual. This analysis estimates a 1349 Mb genome, with 0.249% heterozygosity. The left-hand peak of k-mers corresponds to k-mers from heterozygous regions of the genome, while the right-hand peak is from homozygous regions. **C)** The habitat of the sampled *S. nebulosa lapponica* specimen; photo by Erlend Ragnhildstveit.

DNA for Oxford Nanopore Technologies (ONT) sequencing was isolated from 10 µl blood using the Nanobind® CBB kit (PacBio), following the manufacturer’s protocol for HMW DNA extraction from nucleated red blood cells. Quality check of the amount, purity and integrity of the isolated DNA was performed using a combination of Qubit BR DNA quantification assay kit (Thermo Fisher), Nanodrop (Thermo Fisher), and Fragment Analyser (DNA HS 50kb large fragment kit, Agilent Tech.).

### Library preparation and sequencing for *de novo* assembly

To reduce viscosity, gDNA was homogenized with the DNAFluid+ kit on a Megaruptor 3 (Diagenode), following the manufacturer’s instructions. 5 µg of purified HMW DNA was pipette-sheared into an average fragment length of 36kb using the established PacBio SMRTbell® Prep Kit 3.0 shearing protocol (PacBio Inc.) developed on the Hamilton MicroLab Prep system (Hamilton Nordic AB). The higher input amount (5 vs 3 µg) relative to the original PacBio protocol is what determines the sheared fragment lengths, and in this case tailored for the 30kb ONT library preparation. The sheared DNA was concentrated using SMRTbell capture beads (PacBio Inc) and eluted in 80µl EB buffer before continuing into Oxford Nanopore technology library preparation using the Ligation Sequencing Kit V14 (ONT) following the protocol for Human Variation Sequencing From 30kb Extracted Cell Line Samples (SQK-LSK114), expect changes mentioned above regarding the DNA fragmentation step.

We sequenced the final library on two PromethION R10 flowcells (FLO-PRO114M), with five loadings of 300ng each time on an inhouse P2i system (ONT).

A Hi-C library was prepared starting from approx 20-25µl ethanol-preserved blood using the Arima High Coverage Hi-C kit (Arima Genomics), following the manufacturer’s user guide for nucleated blood samples (Document Part Number: A160177 v01). Final library quality was checked using the QC instrumentation mentioned above for DNA and quantified using a Kapa Library quantification kit for Illumina (Roche LifeScience) on a Lightcycler 96 qPCR instrument (Roche LifeScience). The library was sequenced with other Hi-C libraries on an Illumina NovaSeq X instrument (lllumina Inc) with 2*150 bp paired end mode at the Norwegian Sequencing Centre (https://www.sequencing.uio.no), targeting >50 X coverage.

### Genome assembly and curation, annotation and evaluation

A full list of relevant software tools and versions is presented in Table 1. We assembled the genome using a pre-release version of the EBP-Nor genome assembly pipeline (https://github.com/ebp-nor/GenomeAssembly). KMC (Kokot et al., 2017) was used to count k-mers of size 32 in the ONT reads, excluding k-mers occurring more than 10,000 times. GenomeScope (Ranallo-Benavidez et al., 2020) was run on the k-mer histogram output from KMC and used to estimate genome size, heterozygosity and repetitiveness.

**Table 1.**
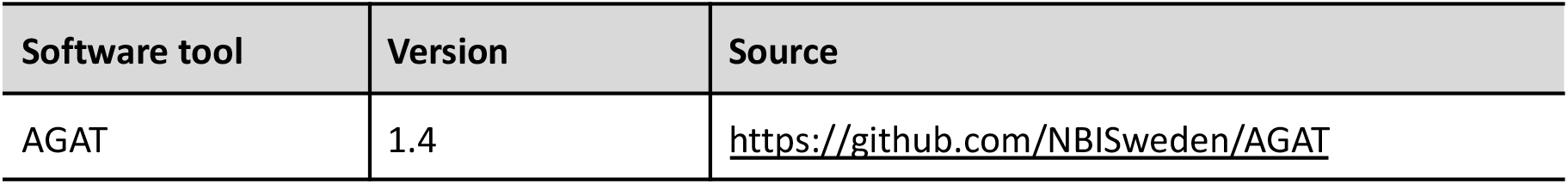
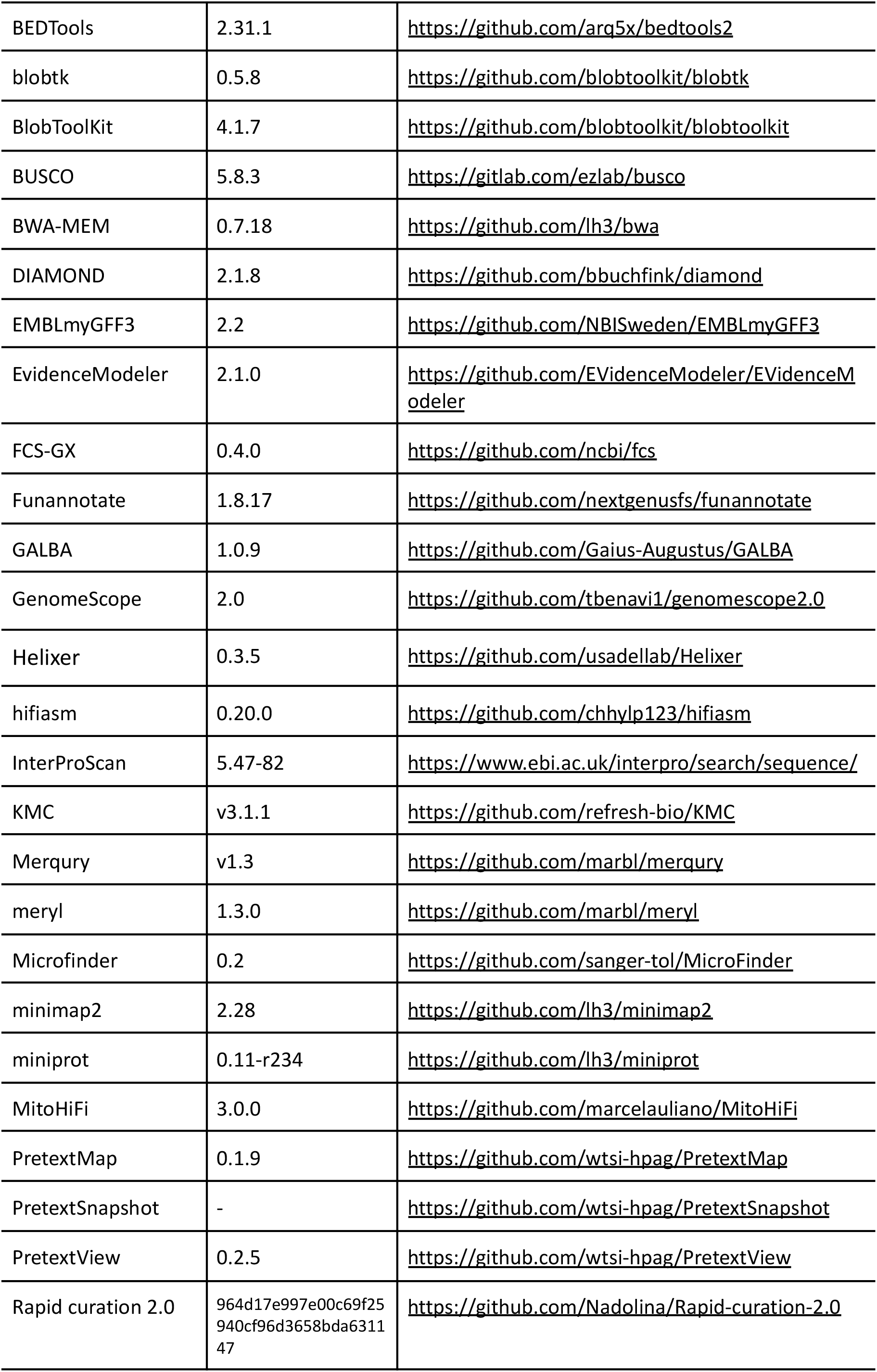
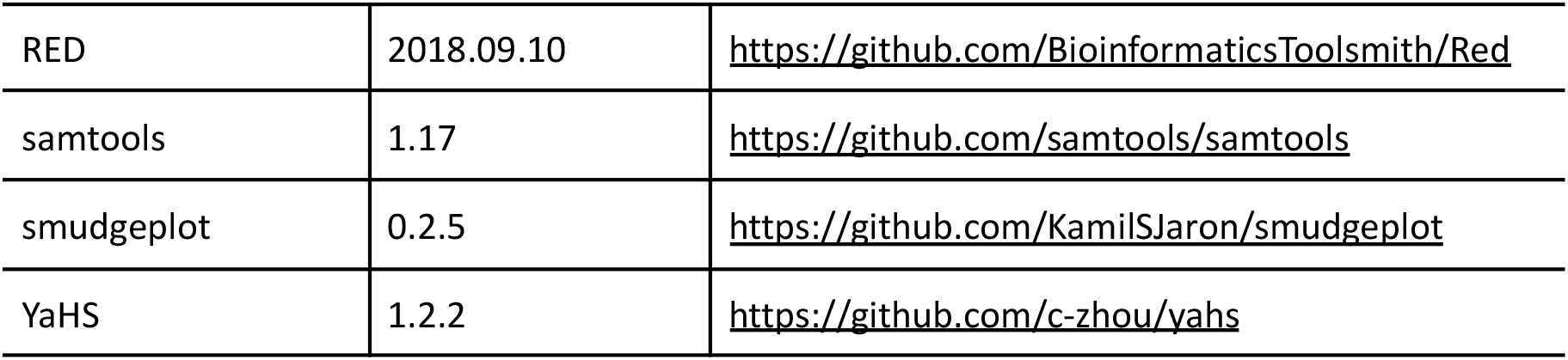
Software tools: versions and sources.

The ONT reads were assembled using hifiasm (Cheng et al., 2021) with Hi-C integration, resulting in two pseudo-haplotype-resolved assemblies: pseudo-haplotype one (hap1) and pseudo-haplotype two (hap2). Meryl (Rhie et al., 2020) was used to identify k-mers unique to each assembly. These k-mers were used to generate two filtered Hi-C read sets: one excluding reads containing k-mers unique to hap1, and one excluding reads containing k-mers unique to hap2. The k-mer-filtered Hi-C reads were aligned to each assembly using BWA-MEM (Li, 2013) with the -5SPM options. Alignments were name-sorted with samtools (Li et al., 2009), after which samtools fixmate was used to remove unmapped reads and secondary alignments and to add mate scores. Duplicates were then removed using samtools markdup. The resulting BAM files were used for scaffolding with YaHS (Zhou et al., 2023) using default options.

The scaffolded assemblies were screened for contamination using FCS-GX (Astashyn et al., 2024), and contaminated sequences were removed. Merqury (Rhie et al., 2020) was used to assess assembly completeness and quality by comparing each assembly to the k-mer content of the Hi-C reads and ONT reads. Assemblies were manually curated in PretextView using Rapid Curation 2.0. Hi-C contact maps were generated by mapping Hi-C reads to the assemblies with BWA-MEM (Li, 2013), using the same parameters as for scaffolding. Contact maps were produced with PretextMap and visualized with PretextSnapshot. Chromosomes, including sex chromosomes, were identified by inspecting the Hi-C contact maps.

A second curation pass focused on putative dot microchromosomes and used assembly scaffolds of 20 Mb or shorter. MicroFinder (Mathers et al., 2026) was used to map conserved dot microchromosome-associated proteins to these scaffolds. The resulting coordinates were used to generate a MicroFinder locus-density BED track for visualization in PretextView. Candidate dot microchromosomes were manually inspected, and curated scaffolds were reintegrated into the assemblies before a final curation pass in PretextView. MitoHiFi (Uliano-Silva et al., 2023) was used to identify mitochondrial sequences in the final assemblies and read datasets.

We annotated the curated genome assemblies using a pre-release version of the EBP-Nor genome annotation pipeline (https://github.com/ebp-nor/GenomeAnnotation). AGAT (https://zenodo.org/record/7255559) scripts agat_sp_keep_longest_isoform.pl and agat_sp_extract_sequences.pl were applied to the *Gallus gallus* bGalGal1.mat.broiler.GRCg7b genome assembly and annotation to generate a reference protein set containing the longest translated isoform from each gene. These proteins were aligned to the curated assemblies using miniprot (Li, 2023). Proteins from the UniProtKB/Swiss-Prot release 2025_03 (UniProt Consortium, 2023) and the Aves subset of OrthoDB v12 (Tegenfeldt et al., 2025) were aligned separately to each assembly.

Repetitive regions in the assemblies were identified using RED (Girgis, 2015), run through redmask (https://github.com/nextgenusfs/redmask), and masked before ab initio gene prediction. GALBA (Brůna et al., 2023; Buchfink et al., 2015; Hoff & Stanke, 2019; Li, 2023; Stanke et al., 2006) was run in miniprot mode using the *Gallus gallus* protein set to generate predicted gene models on the masked assemblies. Helixer (Holst et al., 2025) was run using the vertebrate-specific model vertebrate_v0.3_m_0080.

The funannotate-runEVM.py script from Funannotate was used to run EvidenceModeler (Haas et al., 2008) on the alignments of chicken proteins, UniProtKB/Swiss-Prot proteins, Aves proteins and the predicted genes from GALBA and Helixer. The resulting predicted proteins were compared to the protein repeats that Funannotate distributes using DIAMOND blastp and the predicted genes were filtered based on this comparison using AGAT. The filtered proteins were compared to the UniProtKB/Swiss-Prot release 2025_03 using DIAMOND (Buchfink et al., 2015) blastp to find gene names and InterProScan (Jones et al., 2014) was used to discover functional domains. AGATs agat_sp_manage_functional_annotation.pl was used to attach the gene names and functional annotations to the predicted genes. EMBLmyGFF3 (Norling et al., 2018) was used to combine the fasta files and GFF3 files into a EMBL format for submission to the European Nucleotide Archive.

Genome evaluation was performed using the EBP-Nor genome evaluation pipeline (https://github.com/ebp-nor/GenomeEvaluation). Merqury (Rhie et al., 2020) was used to assess assembly completeness and consensus quality by comparing each assembly with the k-mer content of the Hi-C reads. BUSCO (Manni et al., 2021) was used to assess gene-space completeness using the Aves lineage dataset. Gfastats (Formenti et al., 2022) was used to calculate assembly statistics. Assembly statistics and taxonomic assignments were visualized using BlobToolKit and BlobTools2 (Laetsch & Blaxter, 2017), together with blobtk. Hi-C contact maps were generated by mapping the reads with BWA-MEM using the same approach as described above, followed by PretextMap and PretextSnapshot.

To characterize sequence differences between hap1 and hap2, homologous autosomes were aligned using minimap2 (Li, 2018). The resulting alignments were processed using the minimap2-distributed paftools.js script to quantify single-nucleotide differences and insertions and deletions.

To compare chromosome structure among *Strix* assemblies, we included bStrNeb1_p1.2 (*Strix nebulosa nebulosa*; GenBank accession GCA_051313305.1), bStrAlu1.hap1 and bStrAlu1.hap2 (*S. aluco*; GenBank accessions GCA_031877795.1 and GCA_031877785.1, respectively), and bStrUra1 (*S. uralensis*; GenBank accession GCA_047716275.1; (Chrysostomakis et al., 2025). For consistency, these assemblies are hereafter referred to as bStrNeb1.1.pri, bStrAlu1.1.hap1, bStrAlu1.1.hap2, and bStrUra1.1.pri, respectively. Pairwise whole-genome alignments were generated by aligning each assembly to bStrNeb3.1.hap1 (*S. n. lapponica*) using minimap2 (Li, 2018) with the asm10 preset. Alignments ≥50 kb with mapping quality 60 were retained and visualized in R using gggenomes (Hackl et al., 2024). MicroFinder (Mathers et al., 2026) was run on all assemblies, and retained hits to conserved dot chromosome-associated genes were included in the visualization. Chromosomes and chromosome-associated scaffolds were ordered and oriented relative to bStrNeb3.1.hap1 based on cumulative alignment support. Where multiple sequences corresponded to the same reference chromosome, they were ordered by inferred position along that chromosome to reduce unnecessary crossing of synteny links.

The text in Methods and parts of Results is based on a template we use for all the species we publish in the EBP-Nor project.

## Results

### *De novo* genome assembly and annotation

The genome of the female *Strix nebulosa lapponica* (Figure 1B) had an estimated genome size of 1.35 Gb, with 0.249% heterozygosity and a bimodal distribution based on the k-mer spectrum. A total of 32-fold coverage in R10 Oxford Nanopore reads and 52-fold coverage in Arima Hi-C reads resulted in two pseudo-haplotype-separated assemblies. The final assemblies have total lengths of 1554 Mb and 1242 Mb (Table 2 and Figure 2), respectively. Pseudo-haplotypes one (hap1) and two (hap2) have scaffold N50 size of 56.2 Mb and 88.2 Mb, respectively, and contig N50 of 18.8 Mb and 20.2 Mb, respectively (Table 2 and Figure 2). Both pseudo-haplotypes contained 40 autosomes. For convenience, the W and Z sex chromosomes were included in hap1. Chromosome numbering was assigned in descending order of chromosome length.

**Table 2:**
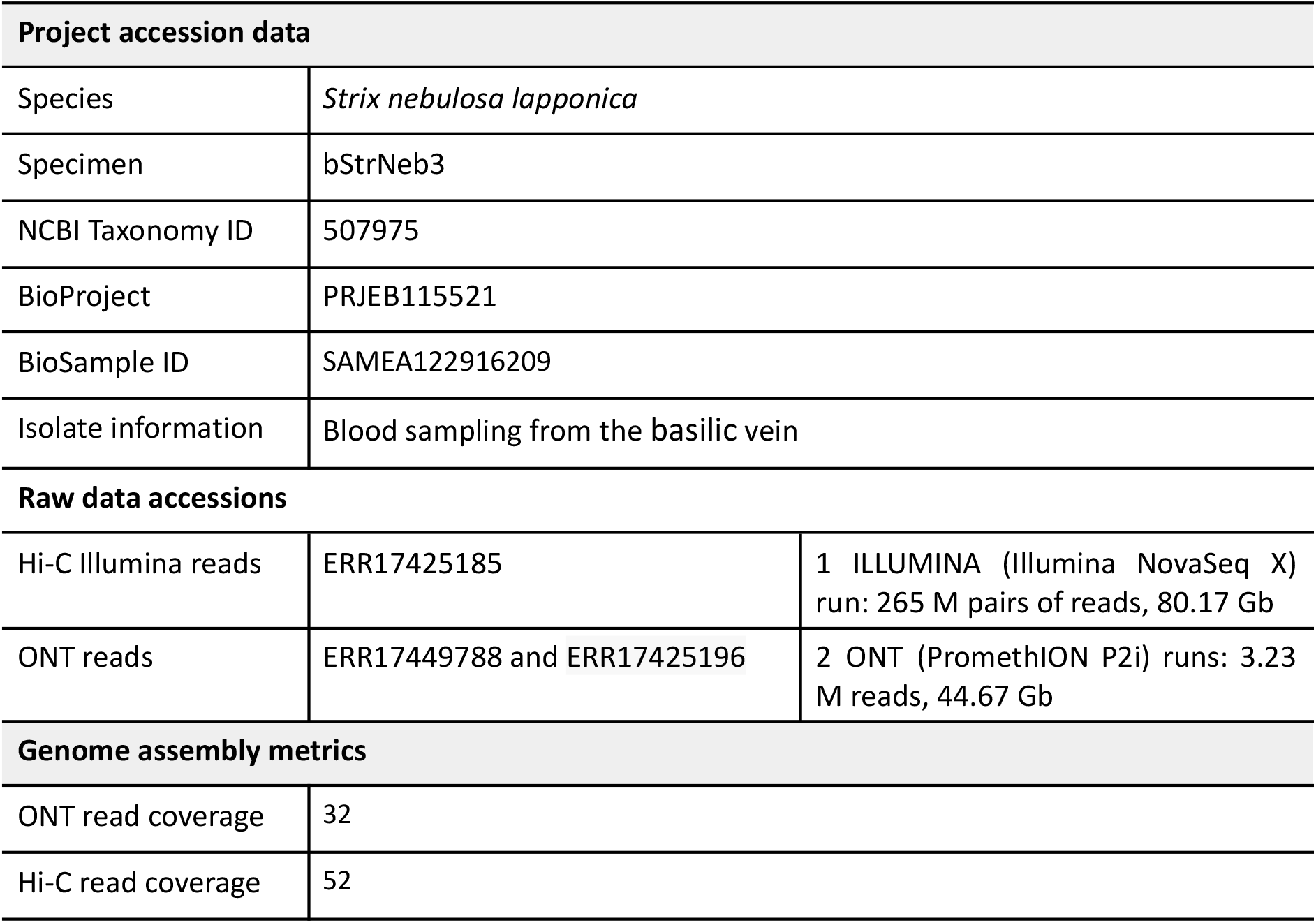
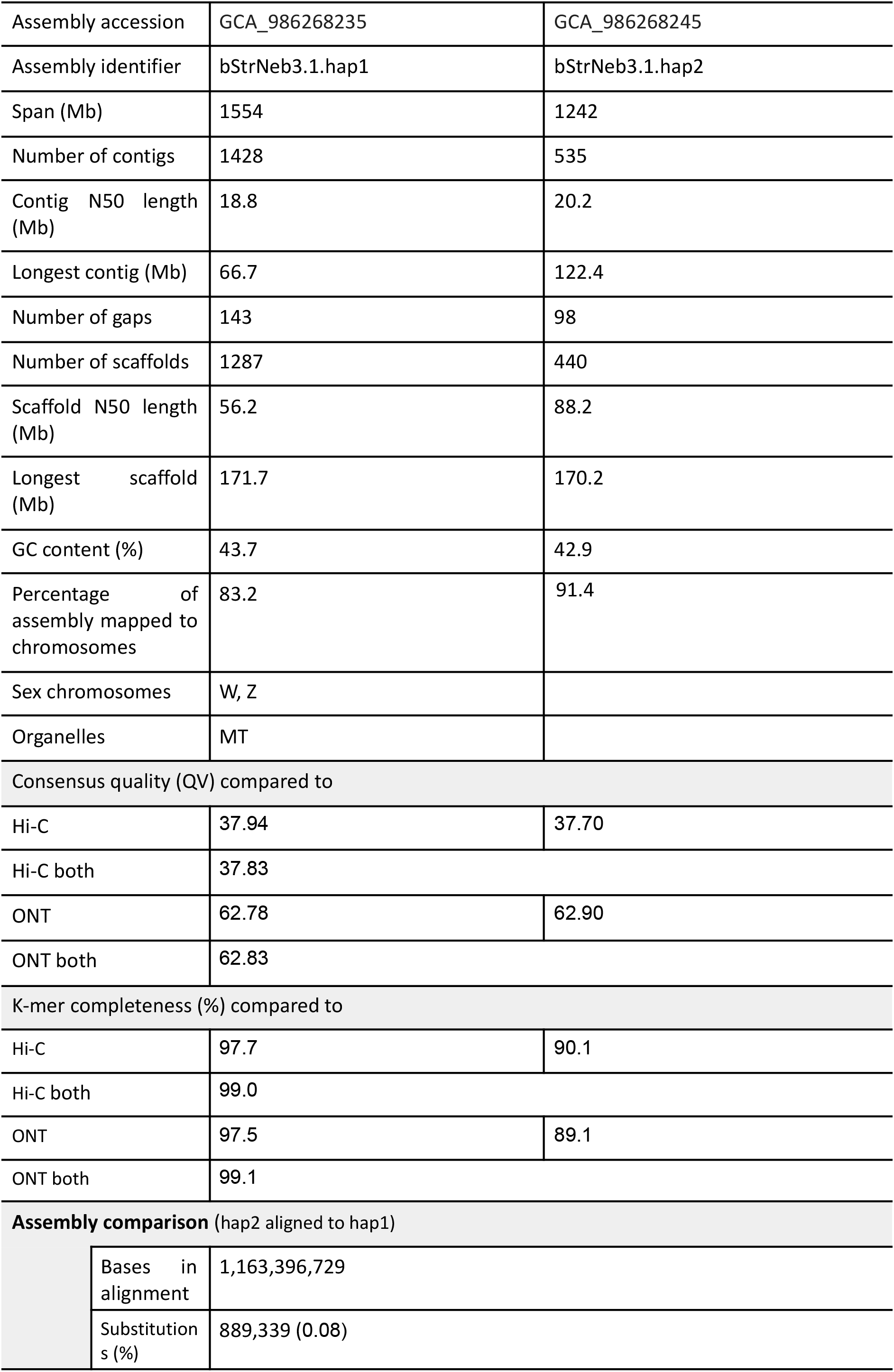
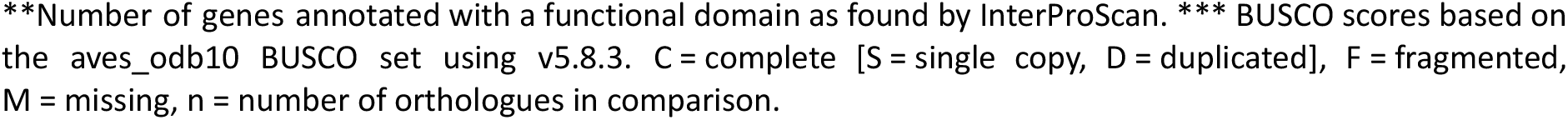
Genome data for *Strix nebulosa lapponica*.

| Project accession data |  |  |
| --- | --- | --- |
| Species | *Strix nebulosa lapponica* |  |
| Specimen | bStrNeb3 |  |
| NCBI Taxonomy ID | 507975 |  |
| BioProject | PRJEB115521 |  |
| BioSample ID | SAMEA122916209 |  |
| Isolate information | Blood sampling from the basilic vein |  |
| Raw data accessions |  |  |
| Hi-C Illumina reads | ERR17425185 | 1 ILLUMINA (Illumina NovaSeq X) run: 265 M pairs of reads, 80.17 Gb |
| ONT reads | ERR17449788 and ERR17425196 | 2 ONT (PromethION P2i) runs: 3.23 M reads, 44.67 Gb |
| Genome assembly metrics |  |  |
| ONT read coverage | 32 |  |
| Hi-C read coverage | 52 |  |
| Assembly accession | GCA_986268235 | GCA_986268245 |
| Assembly identifier | bStrNeb3.1.hap1 | bStrNeb3.1.hap2 |
| Span (Mb) | 1554 | 1242 |
| Number of contigs | 1428 | 535 |
| Contig N50 length (Mb) | 18.8 | 20.2 |
| Longest contig (Mb) | 66.7 | 122.4 |
| Number of gaps | 143 | 98 |
| Number of scaffolds | 1287 | 440 |
| Scaffold N50 length (Mb) | 56.2 | 88.2 |
| Longest scaffold (Mb) | 171.7 | 170.2 |
| GC content (%) | 43.7 | 42.9 |
| Percentage of assembly mapped to chromosomes | 83.2 | 91.4 |
| Sex chromosomes | W, Z |  |
| Organelles | MT |  |
| Consensus quality (QV) compared to |  |  |
| Hi-C | 37.94 | 37.70 |
| Hi-C both | 37.83 |  |
| ONT | 62.78 | 62.90 |
| ONT both | 62.83 |  |
| K-mer completeness (%) compared to |  |  |
| Hi-C | 97.7 | 90.1 |
| Hi-C both | 99.0 |  |
| ONT | 97.5 | 89.1 |
| ONT both | 99.1 |  |
| Assembly comparison (hap2 aligned to hap1) |  |  |
|  | Bases in alignment | 1,163,396,729 |
|  | Substitutions (%) | 889,339 (0.08) |

\*\*Number of genes annotated with a functional domain as found by InterProScan. \*\*\* BUSCO scores based on the aves\_odb10 BUSCO set using v5.8.3. C = complete [S = single copy, D = duplicated], F = fragmented, M = missing, n = number of orthologues in comparison.

**Figure 2:**
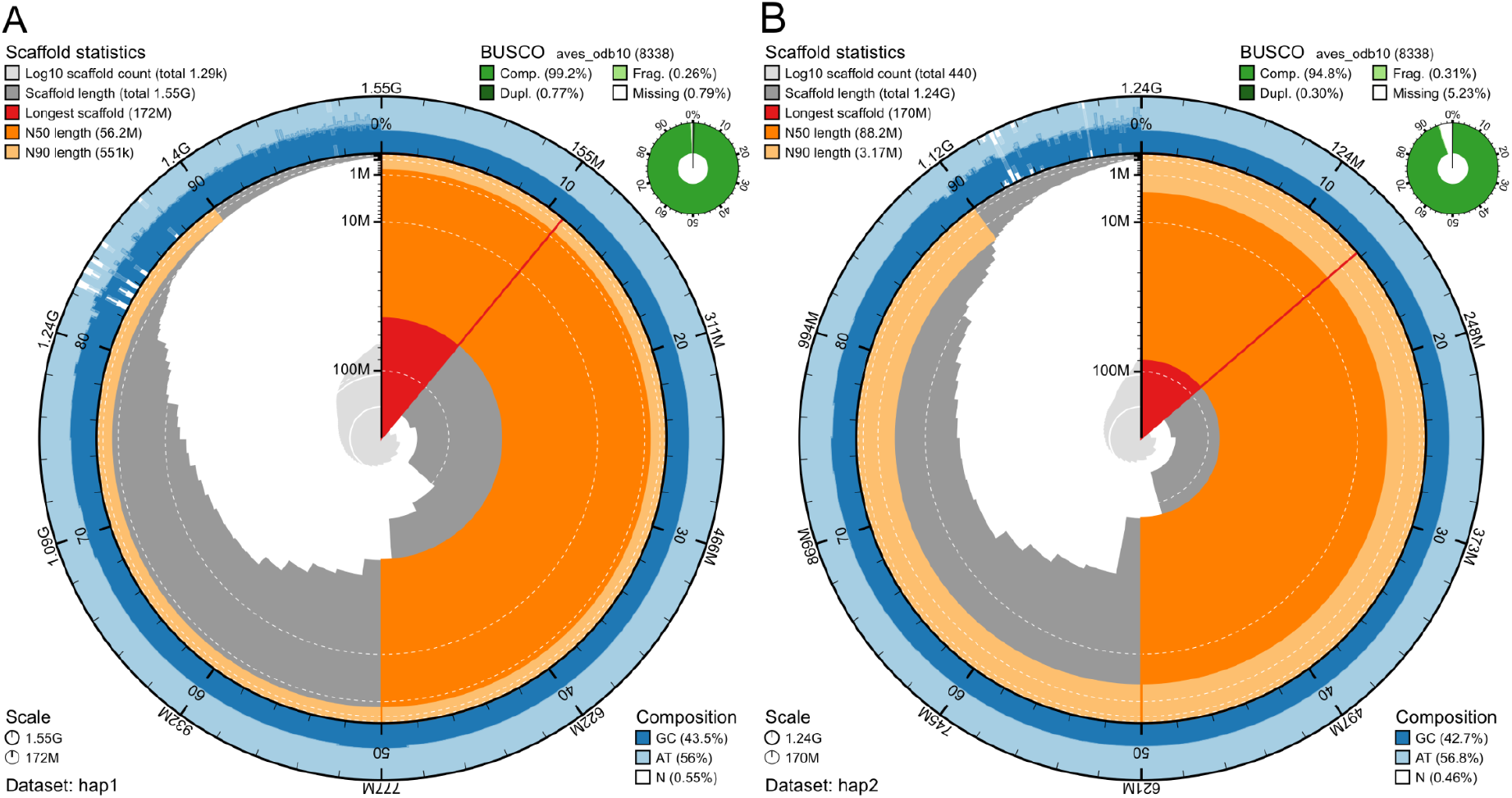
Metrics of the genome assemblies of *Strix nebulosa lapponica* hap1 and hap2. **A)** Hap1. **B)** Hap2. The BlobToolKit Snailplots show N50 metrics and BUSCO gene completeness. The two outermost bands of the circle signify GC versus AT composition at 0.1% intervals, with mean, maximum, and minimum. The third outermost shows the N90 scaffold length, while the fourth is N50 scaffold length. The line from middle to second outermost band shows the size of the largest scaffold. All the scaffolds are arranged in a clockwise manner from largest to smallest, and shown in darker gray with white lines at different orders of magnitude, shown as a scale on the radius. The light gray shows the cumulative scaffold count. The scale inset in the lower left corner shows the total amount of sequence in the whole circle, and the fraction of the circle encompassed in the largest scaffold.

When compared to a k-mer database of the Hi-C reads, hap1 had a k-mer completeness of 97.7%, hap2 of 90.1%, and combined they have a completeness of 99.0%. Further, hap1 had an assembly consensus quality value (QV) of 37.94 and hap2 of 37.70, where a QV of 40 corresponds to one error every 10,000 bp, or 99.99% accuracy compared to a k-mer database of the Hi-C reads (Table 2). When comparing the two pseudo-haplotypes using minimap2, there are 889,339 SNP differences (0.08% of the aligned sequence), 119,654 deletions in hap2 compared to hap1 ranging from 1 bp to more than 1000 bp and 118,773 insertions from 1 bp to more than 1000 bp in size. A total of 18493 and 17279 protein-coding genes were annotated in hap1 and hap2, respectively (Table 2).

The Hi-C contact maps show clear separation of chromosomes into homologous sets (Supplementary Figure 1).

To compare chromosome structure among *Strix* assemblies, we aligned bStrNeb1.1.pri (*S. nebulosa nebulosa*), bStrAlu1.1.hap1 (*S. aluco*), and bStrUra1.1.pri (*S. uralensis*) to the bStrNeb3.1.hap1 *S. nebulosa lapponica* assembly using minimap2. Alignments were visualized with gggenomes after ordering and orienting chromosomes relative to bStrNeb3.1.hap1 (Figure 3A, B). Only alignments ≥50 kb with mapping quality 60 were retained.

**Figure 3.**
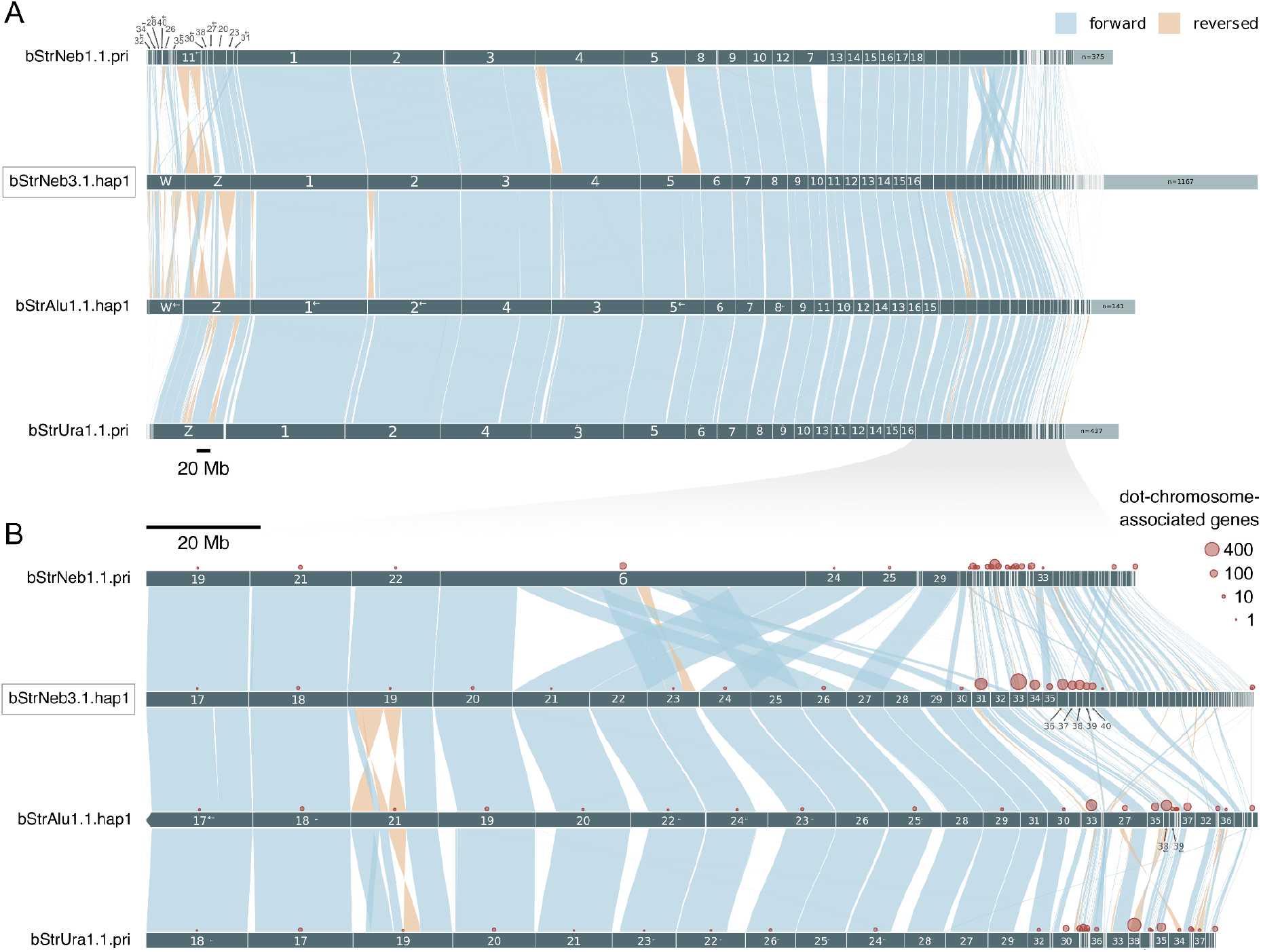
Chromosome-scale synteny across four owl genome assemblies. Assemblies are shown from top to bottom: bStrNeb1.1.pri (*Strix nebulosa nebulosa*), our bStrNeb3.1.hap1 (*S. n. lapponica*), bStrAlu1.1.hap1 (*S. aluco*), and bStrUra1.1.pri (*S. uralensis*). Chromosomes and chromosome-associated scaffolds were ordered and oriented relative to bStrNeb3.1.hap1. When a source chromosome matched multiple sequences, these were arranged by inferred position to minimize crossing links. Blue and orange links indicate forward and reverse alignments, respectively, and arrows mark sequences reversed from their original orientation. Only alignments ≥50 kb with mapping quality 60 are shown. Only scaffolds marked as chromosomes are named. **A)** Full assembly view, including sequences without retained alignments; unconnected non-chromosomal sequences are aggregated. **B)** Enlarged view of source sequences ≤20 Mb and their aligned counterparts. Shading indicates the region enlarged in panel B. Red circles show the number of retained MicroFinder hits to conserved dot chromosome-associated genes, with circle area proportional to hit count.

Macrochromosomes were largely syntenic across the four assemblies and were represented by long, collinear alignment blocks, though several reverse-orientation alignments were observed, particularly near chromosome ends. Extensive fragmented alignments were observed for the Z and W chromosomes and for the smallest microchromosomes. Chromosome 6 in bStrNeb1.1.pri aligned to several distinct microchromosomes in bStrNeb3.1.hap1, bStrAlu1.1.hap1, and bStrUra1.1.pri. The most parsimonious explanation is that chromosome 6 in bStrNeb1.1.pri is a misassembly joining multiple microchromosomes.

Across the six assemblies, MicroFinder detected between 919 and 2,036 of the 2,882 conserved dot chromosome-associated proteins. Recovery was highest in bStrNeb3.1 hap1 and hap2, with 2,036 (70.6%) and 2,000 (69.4%) detected proteins, respectively. Of these hits, 97.6% and 99.8% were located on chromosome-assigned scaffolds (Supplementary Figure 4). In bStrNeb1.1.pri, 1,086 proteins were detected (37.7% of the marker set), of which 27.2% were chromosome-assigned. The bStrAlu1.1 hap1 and hap2 assemblies contained 1,068 and 919 detected proteins (37.1% and 31.9%), respectively, with 89.0% and 70.7% located on chromosome-assigned scaffolds. bStrUra1.1.pri contained 1,492 detected proteins (51.8%), of which 56.4% were chromosome-assigned.

## Discussion

Here, we generated a chromosome-level genome assembly for the Eurasian great grey owl (*Strix nebulosa lapponica*) using ONT R10 long reads and Hi-C data. Compared with the existing assembly from the North American subspecies (*S. n. nebulosa*), the new assembly provides substantially more complete representations of the sex chromosomes and the smallest microchromosomes (Figure 3A, B).

*Strix* assemblies show extensive synteny across the macrochromosomes, with most differences restricted to the sex chromosomes and the smallest microchromosomes (Figure 3A, B). The smallest avian microchromosomes are particularly difficult to sequence and assemble because they are enriched in repeat-associated motifs capable of forming non-B DNA structures, which can substantially reduce or eliminate HiFi coverage (Smeds et al., 2025). ONT-only or ONT-supported assemblies have recovered dot microchromosome sequences that were previously missing or fragmented in bird genomes (Formenti et al., 2025; Huang et al., 2023; Luo et al., 2023). Similar patterns of fragmentation and missing sequence were observed in our comparison of ONT- and HiFi-based assembly of the bluethroat (Strand et al., 2026).

The Bird Chromosome Database reports 2n = 80 for *S. nebulosa* but 2n = 82 for *S. uralensis* and *S. aluco* (Degrandi et al., 2020), with 2n = 82 independently confirmed in *S. uralensis* (Chrysostomakis et al., 2025). Conventional avian karyotyping may underestimate chromosome number because the smallest microchromosomes are difficult to distinguish and count reliably (Malinovskaya et al., 2018, 2022). Thus, the reported 2n = 80 for *S. nebulosa* could reflect the omission of one microchromosome pair.

Using the dot microchromosome-associated protein set of Mathers et al. (2026), our ONT-based assembly recovered more markers than other Strix assemblies, consistent with improved assembly of dot microchromosomal regions. However, these scores should not be interpreted as BUSCO-like completeness scores, because the markers are not one-to-one orthologs expected in all assemblies. We used these results, together with Hi-C and haplotype synteny, to guide manual curation. This resulted in 40 pairs of autosomes plus the Z and W chromosomes, supporting 2n = 82 for *S. n. lapponica*.

Beyond improving the resolution of the smallest microchromosomes and adding both sex chromosomes, this assembly provides a genomic resource for studying evolutionary and demographic processes across the Eurasian distribution of the great grey owl. It is particularly relevant in Norway, where the species remains red-listed despite a recent increase in breeding records and a southwestward expansion of its range. The assembly will be central for comparative genomics, population genetic analyses, and long-term monitoring of this dynamic population near the western margin of the species’ Eurasian range. It will also enable future work on population structure of the whole distribution of the species and to assess possible connectivity between *S. n. nebulosa* and *S. n. lapponica*.

## Funding

This project was funded by the Research Council of Norway project 326819 (The Earth Biogenome Project Norway) to KSJ.

## Acknowledgements

This project received data management and infrastructure support from ELIXIR Norway, supported by the Research Council of Norway’s grant 270068, the University of Bergen, the University of Oslo, the Arctic University of Norway in Tromsø, the Norwegian University of Science and Technology, and the Norwegian University of Life Sciences: NMBU. The authors acknowledge support from the National Infrastructure for High Performance Computing and resources provided by Sigma2 as well as Data Storage in Norway (project NN8013K) for computational work. The Norwegian Sequencing Centre generated the sequencing data used in this project (http://sequencing.uio.no). We are grateful to Jon Are Myhrer and Mette Myhrer for providing invaluable help with setting up the nesting platforms.

## Data Availability

Data generated for this study are available under ENA BioProject PRJEB115521. Base-called ONT reads for the great grey owl (ENA BioSample: SAMEA122916209) are deposited in ENA under ERR17449788 and ERR17425196, while Illumina Hi-C sequencing data is deposited in ENA under ERR17425185. Pseudo-haplotype one can be found in ENA at PRJEB115513, while pseudo-haplotype two is PRJEB115514.

Raw ONT data is deposited at NIRD Archive: https://doi.org/10.11582/2026.yo066exg

The genome assemblies and transposon and gene annotations are available at Zenodo: https://doi.org/10.5281/zenodo.20797770

## Notes

https://doi.org/10.5281/zenodo.20797770

### Competing Interest Statement

The authors have declared no competing interest.

