## Supplementary for "A chromosome-level genome assembly of the Eurasian great grey owl, *Strix nebulosa lapponica* (Thunberg 1798)"

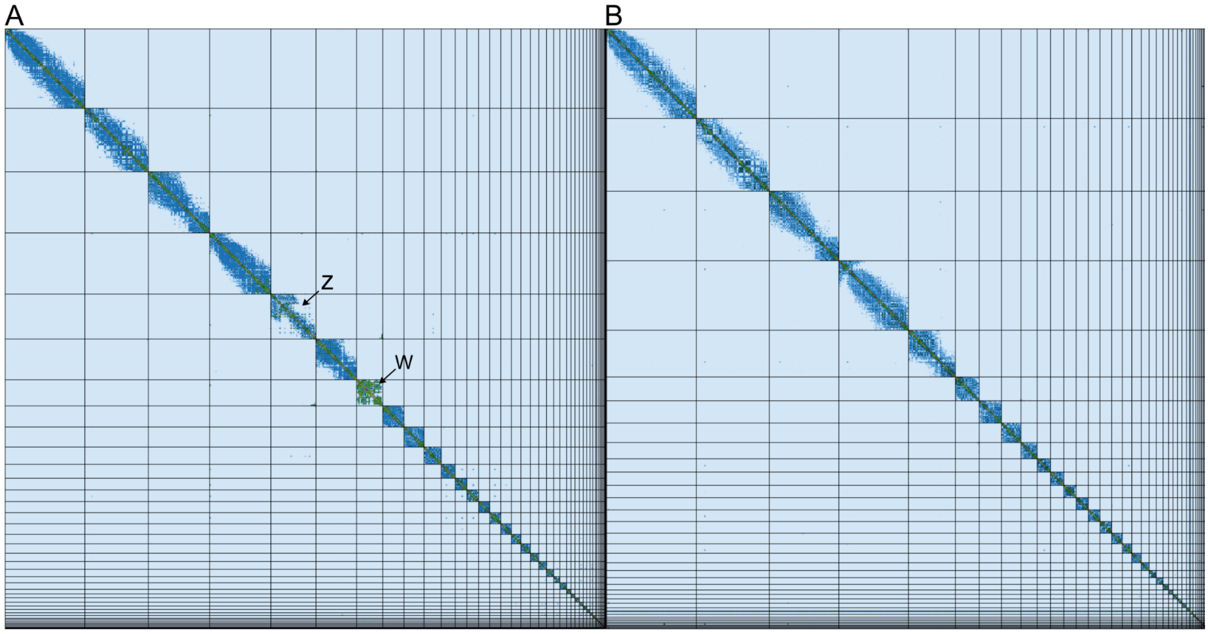


**Supplementary Figure 1: Hi-C contact maps for *Strix nebulosa lapponica* hap1 and hap2.** The contact map displays interaction frequencies between chromosomal regions, where darker shades represent a higher number of Hi-C contacts. The axes correspond to the coordinates along each assembly. The Hi-C contact maps were generated by mapping the Hi-C reads to the pseudo-haplotype genomes using BWA-mem, generating a contact map using PretextMap and visualized using PretextSnapshot. The W and Z sex chromosomes are marked in hap1.

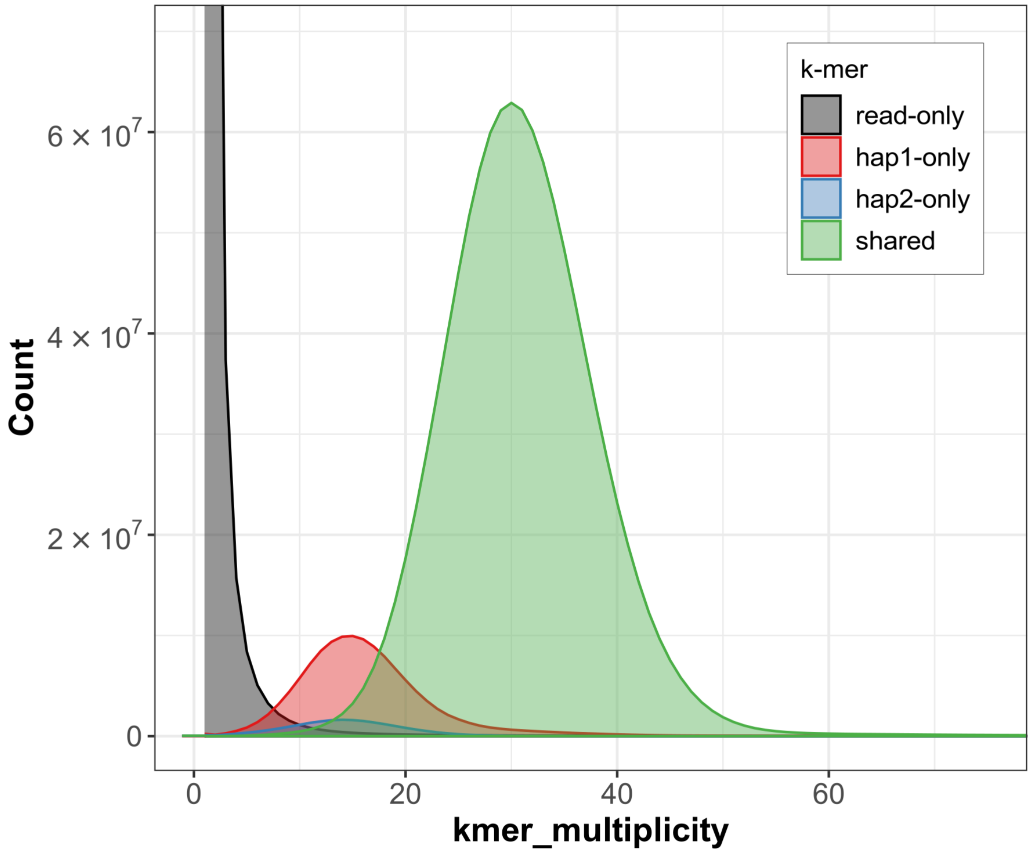

**Supplementary Figure 2: *K*-mer spectra of ONT reads for *Strix nebulosa lapponica*.** Distributions of *k*-mers found only in the reads (black), only in haplotype 1 (red), only in haplotype 2 (blue), or in both haplotypes (green). The y-axis represents the number of unique *k*-mers, while the x-axis represents the *k*-mer multiplicity (how often the *k*-mer is found in the set of ONT reads).

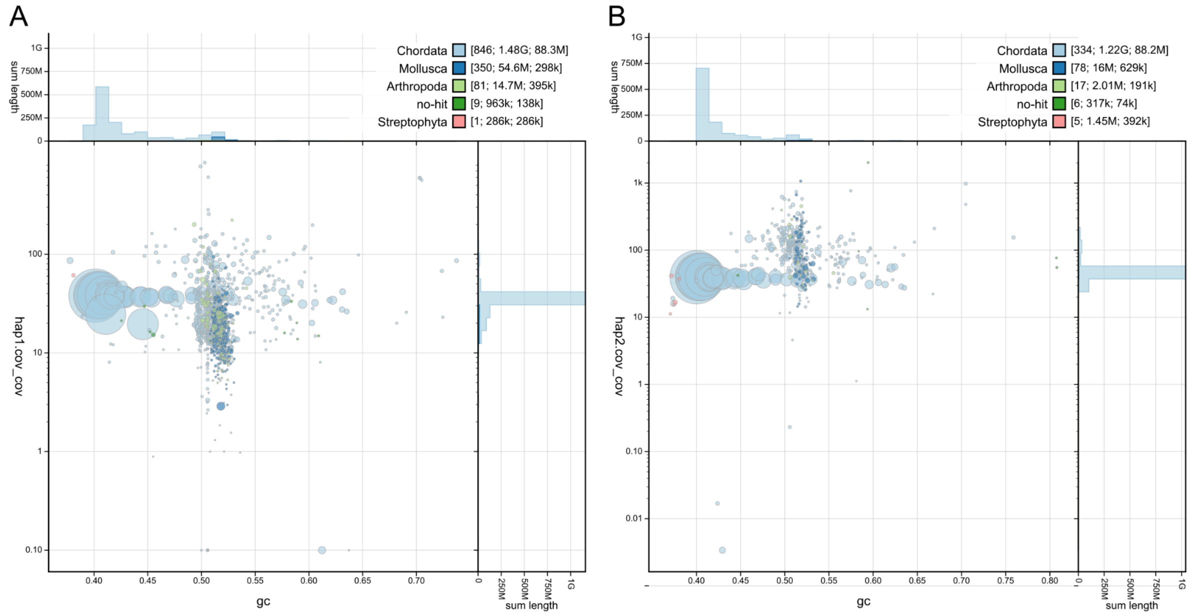


**Supplementary Figure 3: Coverage vs GC plots of *Strix nebulosa lapponica* hap1 and hap2. A)** Hap1 **B)** Hap2. The BlobToolKit Blobplots depicts each scaffold as a dot based on the GC content (%GC, x-axis) and coverage (Y-axis). Size of the dots correspond to scaffold length. Dots are colored based on assigned taxonomy. Histograms of sequence lengths within a certain %GC range or coverage range are depicted on the top and right respectively.

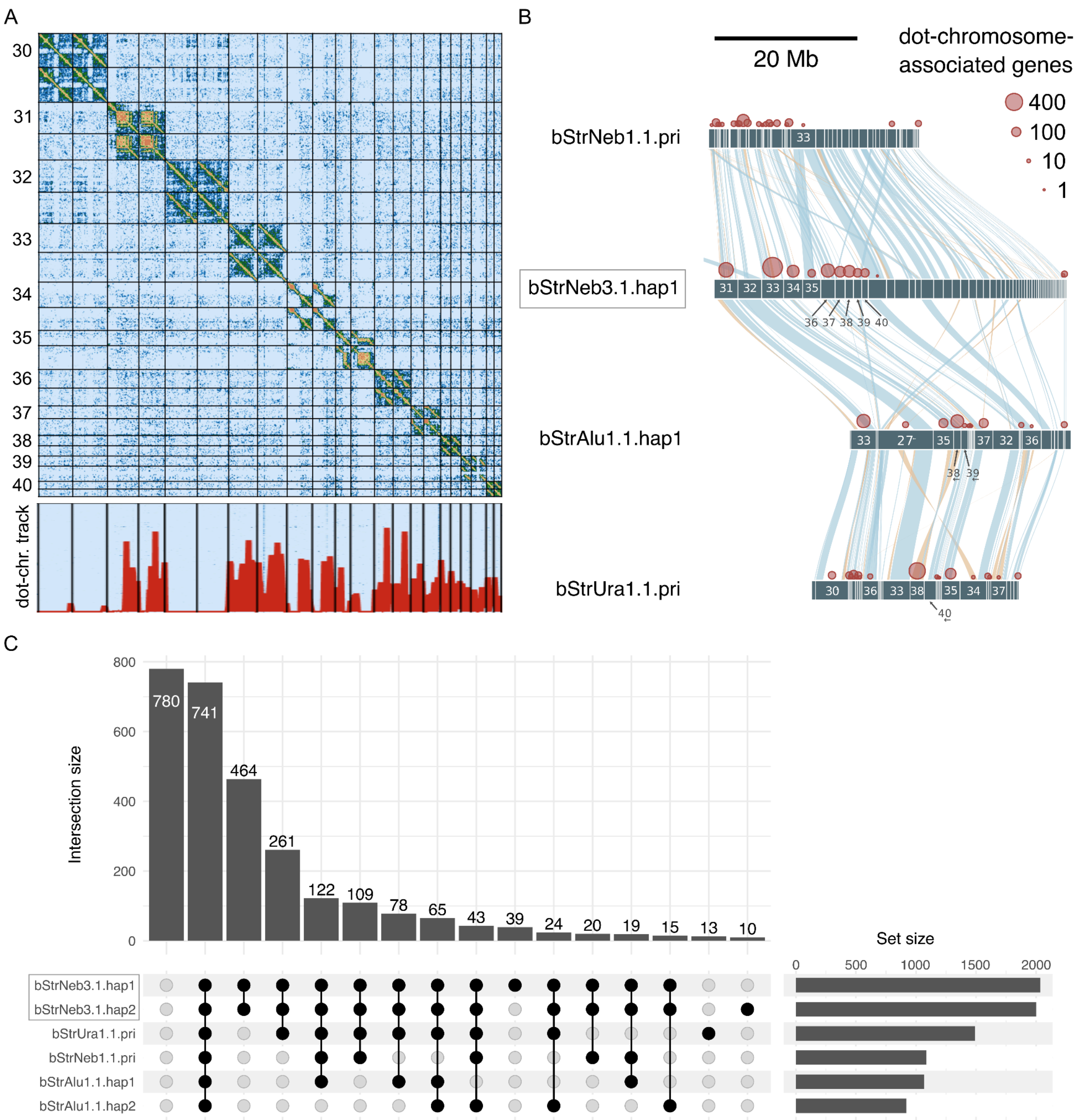


**Supplementary Figure 4. Hi-C support and conserved gene content of the smallest chromosomes. A)** Hi-C contact map of the 11 smallest chromosomes in Strix nebulosa lapponica hap1 and hap2 (hap1 comes first from the top followed by hap2). The MicroFinder locus-density bedGraph track below the contact map shows the distribution of conserved dot chromosome-associated genes. **B)** Enlarged synteny view of the 10 smallest chromosomes and their associated scaffolds across the four Strix assemblies shown in Figure 3. Blue and orange links indicate forward and reverse alignments, respectively. Red circles mark retained MicroFinder hits to conserved dot chromosome-associated genes, with circle area proportional to the number of hits. C) Upset plot showing the overlap in conserved dot-chromosome-associated genes detected by MicroFinder across the six assemblies. Vertical bars indicate the number of genes shared by each exact combination of assemblies marked by connected black dots, while horizontal bars show the total number of genes detected in each assembly.
